# A Low-Cost Markerless motion capture system to automate Functional Gait Assessment: Feasibility Study

**DOI:** 10.1101/2024.12.17.628985

**Authors:** Osman Darici, Chanel Cabak, Jeremy D Wong

## Abstract

Functional gait assessments in older adults have traditionally required manual in-person quantification of clinical measures such as walking speed and step placement. This reliance on individuals trained in motion analysis hinders the frequency with which they are performed, and reduces their generalizability and replicability. To standardize, simplify, and broaden access to gait assessments we here deploy recently-developed open-source tools to produce a low-cost, AI driven markerless motion capture system with custom analysis software for Functional Gait Assessment. Our system uses 3 Cameras, and was validated with a traditional marker-based system on subjects (N=3), showing strong correlation to laboratory-standard measures of step length (*R*^2^=0.98), step width (*R*^2^=0.97), and head speed (*R*^2^=0.95). The markerless system’s FGA reports demonstrated data similar to previous FGA findings in older adult subjects (N=5). Moreover, supplemental standard biomechanical gait measures Step Width, walking Duration, and continuous Gait Speed may be integrated to augment existing FGA. This study demonstrates a proof-of-principle open-source markerless system for analyses of functional gait.

## Introduction

Due to cost and complexity of biomechanical analysis tools such as motion capture, standard clinical human gait tests are typically designed to be easy to perform by a clinician. However, these tests still cost time and attention to administer and may also be sensitive to inter-clinician variability. This is true for the tests such as Functional Gait Assessment (FGA) (Wrisley and Kumar, 2010), Timed Up and Go (Herman et al., 2011), and Edinburgh Visual Gait Score (Aroojis et al., 2021), each of which are different protocols meant to monitor gait for clinical decision making and fall risk prediction. FGA does not require sophisticated equipment but instead demands a sophisticated observer. Ten different gait conditions—such as walking with changes in speed, motion of the head, or while stepping over obstacles—are performed during which task time is recorded via stopwatch and lateral step placements, gait speed changes, response time, and step height over obstacles are visually monitored. Some conditions require subjective judgments such as whether a speed change was “significant” or “minor”. The Timed Up-and-Go test is simpler, requiring total duration of a 3m out-and-back walk, from a seated start to end; yet task time (presumably done visually via stopwatch), and aberrations to gait and balance are only visually monitored. The Edinburgh Visual Gait Score outputs more sophisticated gait metrics such as joint angles, requires (in addition to an attentive clinician) substantial processing time (Aroojis et al., 2021), and still may lack of inter-rater reliability (Ong et al., 2008; Viehweger et al., 2010). Advances in markerless motion tracking leveraging artificial intelligence (AI) may enable low-cost motion capture systems to be utilized to perform these clinical tests and enhance their limitations. Therefore, in this paper we focused on implementing an open-source motion capture system employing pose-detection via AI and computer vision to perform one of these clinical tests: FGA. We aimed to show that a user-friendly, low cost markerless motion system can improve measurement accuracy and therefore reduce inter-clinician variability and also can output several metrics to improve clinical relevance of the test.

An obstacle to motion capture-based gait analysis research is the complexity and cost in analyzing walking behaviour. Motion capture data—the most widely accepted method for analysis of full-body locomotion—requires costly equipment to collect, substantial time to analyze, and skill to convert kinematic data into clinically relevant gait metrics. Dedicated spaces are often used to allow a set of cameras to encircle a subject. Markers placed on limb segments consume substantial time, and cameras must be correctly calibrated to convert 2D images recorded from cameras into 3D positions. Commercial markerless laboratory systems have begun to be used which eliminate the use of markers at comparable accuracy (Lichtwark et al., 2024; Nakano et al., 2020), but remain accessible only to research institutions due to both the operational costs of annual software licensing and complex analysis.

Advances in AI-driven computer vision have led to useful pose estimation and markerless motion capture techniques (Cao et al., 2019; Graving et al., 2019; Guo et al., 2022; Ramesh et al., 2023; Uhlrich et al., 2023) that are open-source, but research is needed to show that these tools can automate clinical assessments. One obstacle to gait assessment automation with open-source tools is the use of multiple cameras and custom algorithms to provide reliable, clinically-relevant metrics. Studies utilizing monocular video for example, do not demonstrate highly-accurate estimates of step width, despite being reliable at providing gait asymmetry estimates stroke patients (Stenum et al., 2024; Wang et al., 2024). Functional gait tests typically use step width as an indicator of decreased stability and elevated fall risk, and step width changes during gait challenges such as eyes-closed or narrow-base-of-support walking is one common clinical assessment approach. To our knowledge it remains to be demonstrated if markerless open-source motion capture technologies can provide reliable gait metrics within a Functional Gait Assessment.

A low cost, user-friendly, markerless motion system may make clinical gait tests more comparable to the broader biomechanics literature. For example, mediolateral foot placement is used in FGA, possibly because it is easier to observe manually, but confounds gait heading and step width together; step width is widely understood to primarily reflect a human’s mediolateral balance control (Bruijn and van Dieën, 2018; Donelan et al., 2001; Kuo, 1999; Redfern and Schumann, 1994; Wang and Srinivasan, 2014; Winter, 2009). Moreover, while total movement duration is typically measured in a clinical setting, continuous measurement of a gait speed trajectory reflects the control of non-steady gait (Brown et al., 2021) and thus applicable to FGA tasks, and which may provide information about an individual’s value placed on energetic cost and movement time (Carlisle and Kuo, 2023). Such outcomes can be made accessible with a markerless motion system, supplemented with automated gait analysis.

In this study, we chose The FreeMoCap Project (Matthis et al., 2022) to measure body segment position, supplemented with custom code for FGA analysis in an effort to determine if such a system could reduce measurement subjectivity and automate assessment. FreeMoCap represents one of the first among a set of open-source 3D computer vision projects capable of integrating pose estimations from multiple cameras into estimated of segment locations with consumer-grade cameras. FGA is an assessment widely used in clinics and which is a versatile test involving complex movement patters, such as turning, speed changes, and obstacle navigation. We utilized 3 consumer-grade web-cameras and compared the accuracy of the system to a standard marker-based motion capture system. We then measured older adults performing FGA and developed custom software to automatically output performance metrics of FGA without any human interaction. Furthermore, we extracted clinically relevant step width and speed trajectories for each condition of FGA which cannot be determined by visual inspection. We therefore tested the prediction that an automated system could reliably perform FGA, automatically output test results and improve clinical relevance by automating scoring based on step length, step width and walking speed.

## Methods

### Ethics

Young (N=3) and older adult subjects (N=5; ages 65-79) provided their written informed consent according to institutional review board procedures.

### Experimental Designs

We performed two experiments. First we established the validity of the opensource markerless motion capture system (FreeMoCap: Matthis et al., 2022) to record clinically-relevant metrics—step width, step length, and walking speed—by comparing markerless foot placement and the resulting step times and durations, to those from an active optical motion capture system commonly used in biomechanical studies (Phasespace Inc.). Next, we performed Functional Gait Assessment on Older adults (Wrisley and Kumar, 2010). We placed the analysis code in a repository for use by practitioners interested in performing their own analysis.

### Methods

The markerless motion capture system used 3 (Jetaku Inc; Amazon) web cameras placed laterally to the walking direction along a 6 meters distance which is used in FGA (Figure 1A). We also marked the hallway to denote a 6 m walkway, with a witdh of 0.3 m. A desktop PC (2018; 3.4 GHz; 4 Cores; 16 GB Ram) was used to record and process data. Total added cost of the cameras and USB extension cables was less than $200 (CAD).

**Figure 1.**
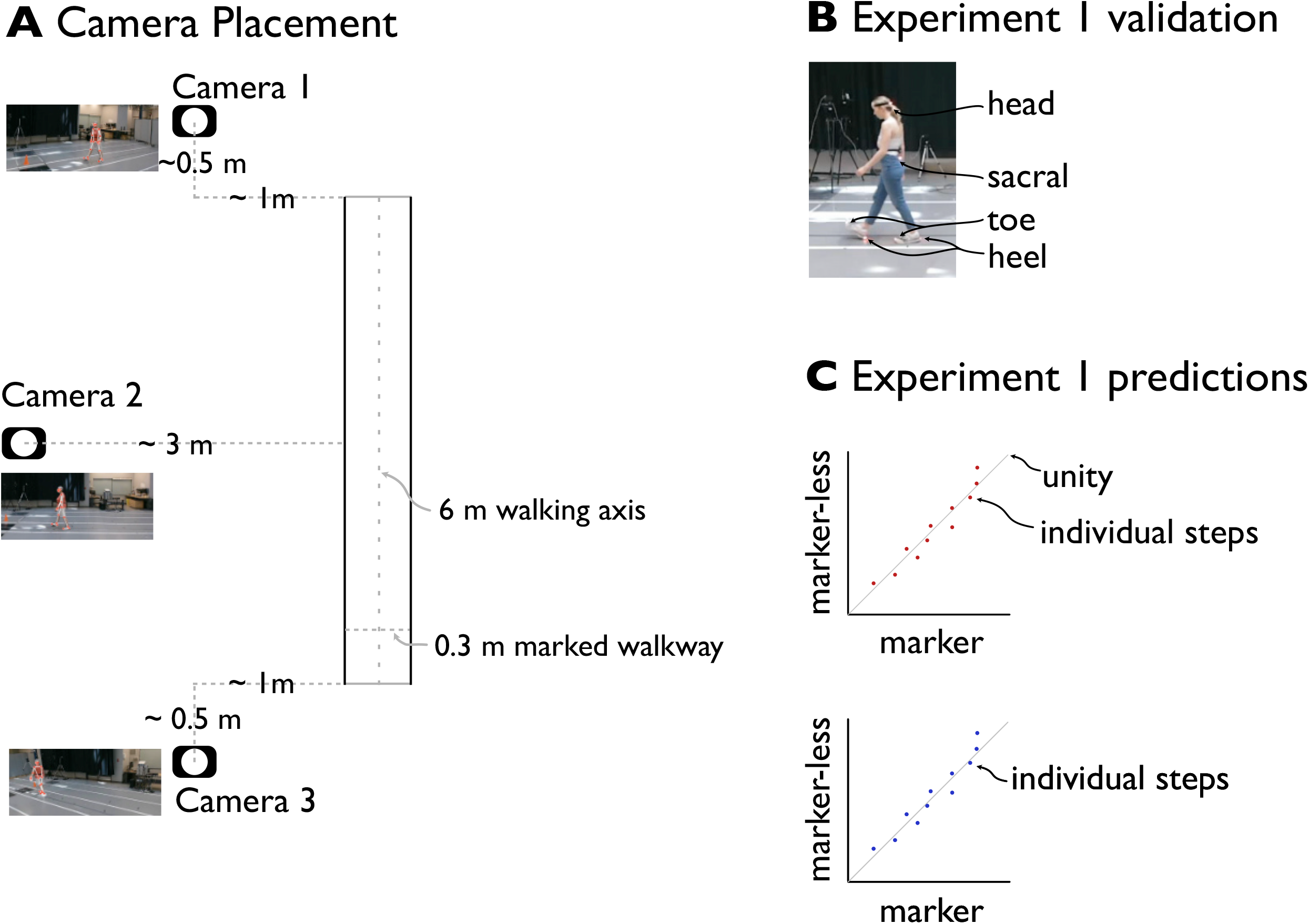
Experimental setup and validation experiment. A: Camera placement, with approximate distances relative to the left edge of the walkway (note: depicted person is of one of the authors). B: Experimental validation of step placements, using markers placed on the feet, sacrum and head. C: Validation of markerless data: step length, width and walking speed.

Experiment sessions required approximately 20 minutes of preparation time to 1) physically set-up cameras, 2) perform calibration and 3) compute capture volume orientation. To calibrate the system, the markerless motion capture system optimally estimates the 3D position of web cameras, and accounts for visual distortion of the camera optics. This is done by simultaneously recording video data from a Charuco board—a single black-and-white calibration image with known checkerboard pattern and dimension. To accommodate a 6m walking area that FGA requires, a large poster (square size: 15.7 cm) was used. Custom software optimizes the 3D position of the cameras to minimize the 3D-positional error of the detected checkerboard patterns (or “bundle adjustment (Karashchuk et al., 2021)”). The software (Matthis et al., 2022) reports the number of full boards detected from each camera, and provides some information about calibration accuracy; practically, we ensured that at least 400 boards were recorded for from each of the web cameras for each of our recording sessions. This calibration process took 2 minutes of Charuco board positioning, and about 10 minutes of compute (no GPU acceleration; CPU processing only on our 2018 machine). To perform capture volume orientation, subjects stood in three locations to define a simple coordinate system: 1) at the beginning of the walkway; 2) at +5m (forward) along the pathway; and 3) at 1 m right of the walkway. This procedure defined the plane of the walking area, allowing rotation of the recorded data. It also allowed us to perform linear correction for calibration bias, as these known magnitudes allowed rescaling data collected along each axis. Typically, FreeMoCap’s calibration procedure resulted in less than 5% error in absolute estimate of walkway length.

During data processing, key-point software makes estimates of the position of joints and other points of interest on the body (Google Mediapipe). The software performs a lowpass filter (25 Hz) to reduce the effect of joint misestimation, motion blur, and other artefacts.

### Experiment 1: Validation

We performed our first experiment to determine the validity of the markerless motion capture system to compute three measures that are necessary and sufficient to automate FGA: step length, step width, and walking speed. To determine the accuracy of computer vision methods, we used marker-based motion capture and collected 6 markers per person to capture feet, torso, and head position of subjects (N=3). We simultaneously recorded walking trials of a 4 m distance with both systems, under two conditions chosen to span the range of empirically-observed step widths: normal preferred step width, and a ‘wide’ condition where subjects were asked to place footsteps along the edges of the 30 cm marked path. We performed three replications of each.

We compared the estimated positions of the two systems by using the heel marker position from the markered system to the foot position from markerless motion capture. We time-aligned the two systems by determining the maximum cross-correlation between the two extracted walking bouts after up-sampling the markerless system (30 Hz) to the markered system’s framerate (240 Hz). We subtracted the initial (T=0) position of each left foot to register the two systems at the beginning of the walking bout.

We computed step length and width as the distance between two consecutive foot placements. We measured head speed as the motion of the head marker (located centrally in the back of the head via headband), and for the markerless system we defined it as the average of the left ear, right ear, and nose keypoints.

### Experiment 2: FGA

Older adult subjects (N=5; mean age: 83 years) were asked to perform a complete functional gait assessment. We performed three repetitions of each task.

### Markerless Setup: lighting and clothing

Accurate detection of body segments is sensitive to visual elements that make it difficult for the Mediapipe algorithm to clearly separate body keypoints from the background. Moreover, illumination—which affects shutter speed and thus motion blur—can affect localization. Optimal algorithm performance was achieved by using contrasting shoes with the ground colour—either white or dark shoes against the grey floor—and similarly contrasting colours for clothing, that fit relatively tight to the body. These adjustments can substantially affect the algorithm’s tracking accuracy. For each subject we ensured that body keypoints during a practice trial were consistently and smoothly detected before proceeding with data collection, and swapped out poorly-detected shoes or clothing if keypoint detection was problematic.

### Data analysis

We developed custom MATLAB code to analyze walking bouts for Functional Gait Assessment, which uses a detailed set of manually-extracted gait features but relies on step placement and walking speed. We cross-correlated (matlab: normxcorr2) each recorded trial with a template of forward footspeed (captured during Experiment 1), and the maximum value was used to compute the walking bout’s start and end timepoint. Foot speed was low-pass filtered at 3 Hz, and regions where foot speed was below the speed threshold of 0.15 m/s were identified as stance regions. The average position of that stance region was taken as the step location. Step timing was computed over the interval between consecutive midstance points.

### Functional Gait Assessment Scores: Automation

FGA (Wrisley and Kumar, 2010) is meant to evaluate a person’s ability to maintain stability while walking under various conditions, and consists of 10 walking conditions (changes in speed, walking with head turns, stepping over obstacles) each of which is scored between 3 (highest score) and 0 (lowest score). For some trials, qualitative changes in behaviour are judged by an observer, which we here automated according to measured quantities. For example, FGA Scores for Condition 2 require that a “Change in Gait Speed” be determined as “significant” for highest score; similarly Conditions 3 and 4 (gait with horizontal and vertical head turns) require that “no change in gait” be detected for highest score (Wrisley and Kumar, 2010). We defined the following quantities relative to a person’s baseline level collected during Condition 1 (normal walking), and note that these values demonstrate proof of principle of assessment automation.

Condition 2: *Gait Speed Change*:

∘ FGA score 3: speed change greater than 125% Condition 1 (normal walking)
∘ FGA score 2: speed change between [110%, 125%] Condition 1
∘ FGA score 1: speed change between [105%, 110%] Condition 1
∘ FGA score 0: no speed increase greater than 105% of Condition 1

Conditions 3 and 4: *Gait with Horizontal/Vertical head turns*:

∘ FGA score 3: maximum speed within [5%] of Condition 1, mediolateral placements as noted in (Wrisley and Kumar, 2010) Condition 1
∘ FGA score 2: speed within 10% of Condition 1, mediolateral foot placements as noted in (Wrisley and Kumar, 2010)
∘ FGA score 1: speed change within [75%, 90%] of Condition 1
∘ FGA score 0: speed change less than [75%] Condition 1

Condition 5: *Gait And Pivot Turn*:

∘ FGA score 3: turning in less than 3 s
∘ FGA score 2: turning (cued verbally at 2m into walk) took longer than 3 s
∘ FGA score 1: turning (cued verbally at 2m into walk) took longer than 4.5 s
∘ FGA score 0: incomplete

Condition 6: *Step Over Obstacle:*

∘ FGA score 3: high obstacle clearance, speed greater than 75% Condition 1
∘ FGA score 2: lower obstacle, speed greater than 75% of Condition 1
∘ FGA score 1: lower obstacle, speed less than 75% of Condition 1
∘ FGA score 0: incomplete

## Results

### Experiment 1

The markerless motion capture system was accurate in its detection of foot placement events (Figure 2A). The Markerless foot positions showed good agreement with the marker-based motion capture during both normal and wide step widths, demonstrating high correlation in step length (*R*^2^=0.98; Figure 2B) and step width for both normal and wide trials (*R*^2^=0.97; Figure 2C).

**Figure 2.**
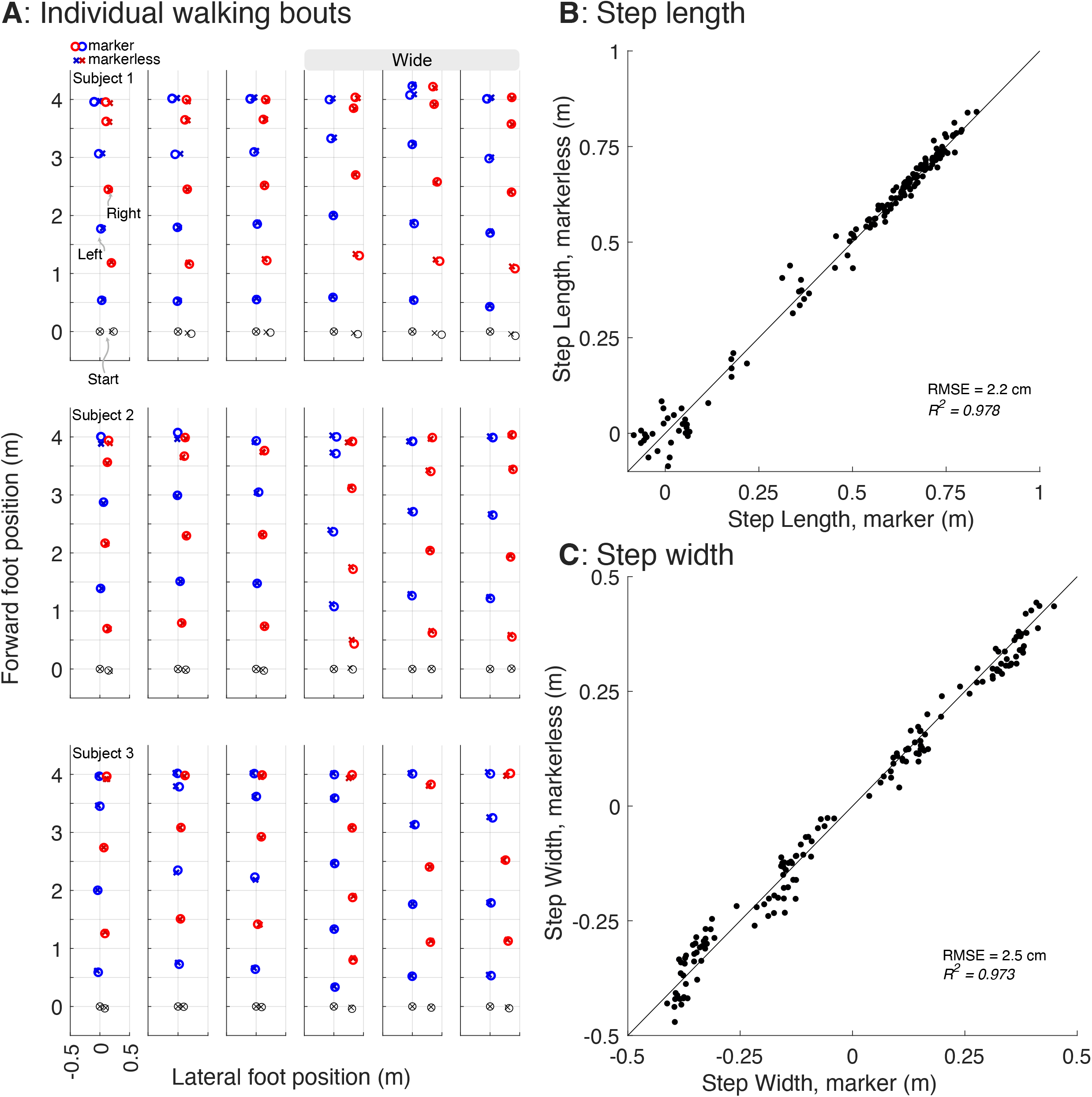
Comparison of markerless system foot placement estimates with traditional marker-based motion capture. A: individual steps of all subjects of both normal and wide walking trials identified via marker (O’s) and markerless motion capture (X’s). B and C: Step length (B) and width (C; negative is left-to-right step width) for each step shown in A.

Head speed measures with the markerless system were also similar to marker-based motion capture. Figure 3 depicts step speed across the normal and wide trials (Figure 3; *R*^2^ = 0.91 to 0.98; mean = 0.95).

**Figure 3.**
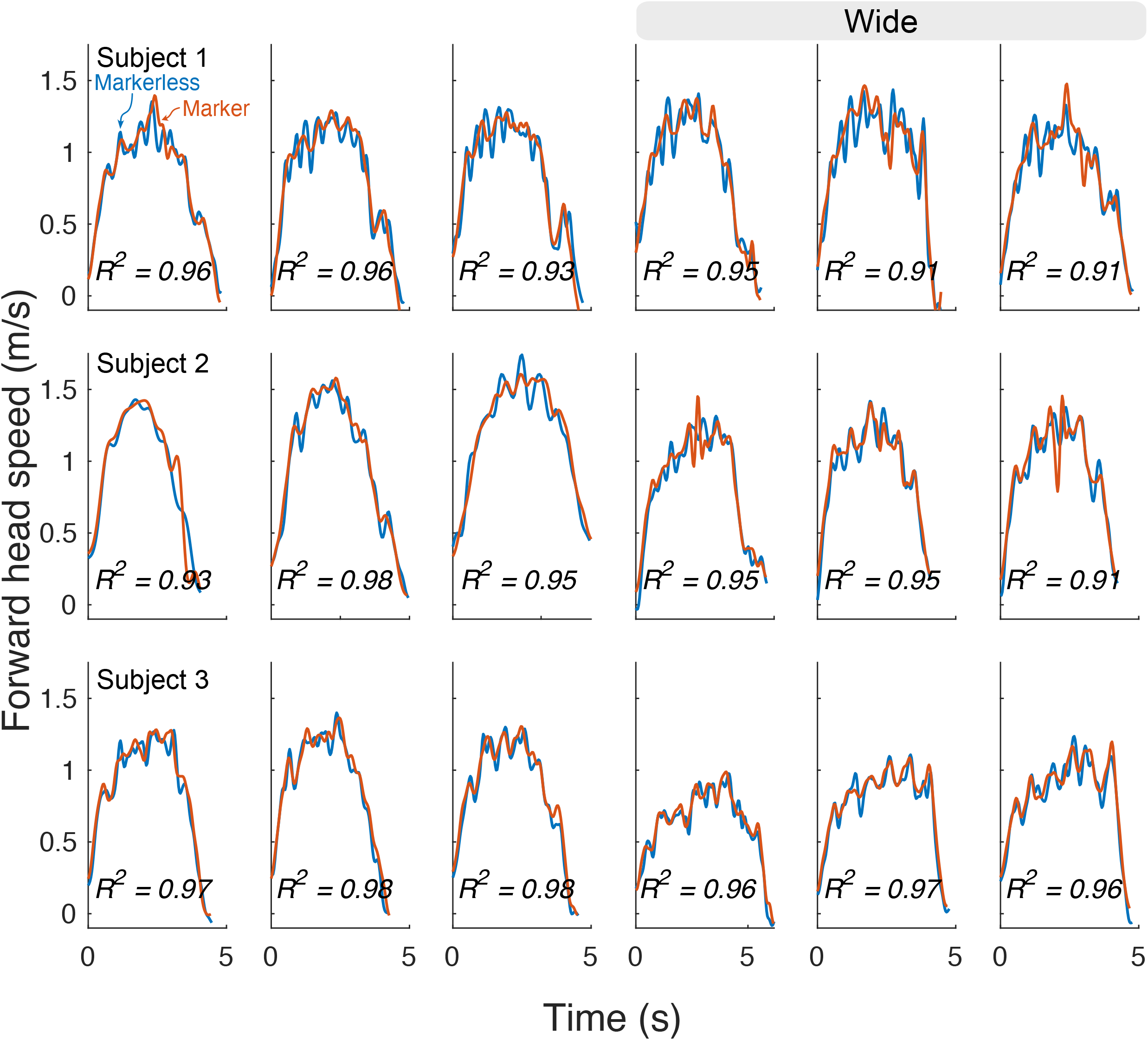
Head speed measured during walking trials, by markerless (blue) and marker-based (red) methods.

### Experiment 2

In Experiment 2, we employed the Functional Gait Assessment (FGA) for our older adult subjects, which resulted in quantification of individual footsteps (Figure 4A) for each condition. Head speed trajectories (Figure 4A, bottom) were sufficient to show deviations in walking speed (forward speed) per condition, relative to nominal preferred walking (Trial 1).

**Figure 4.**
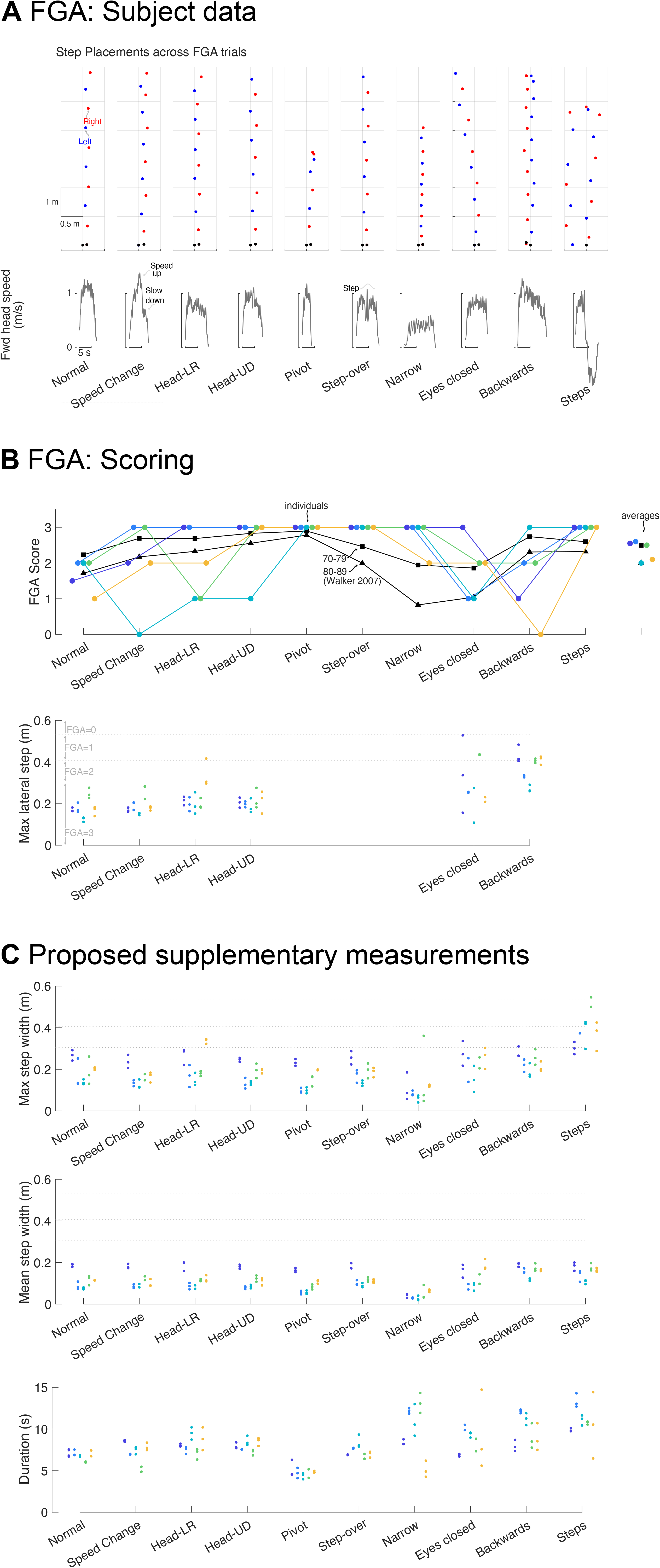
Functional Gait Assessment. A: Representative subject data, step placements for left (blue) and right (red) feet for all 10 conditions. Head-LR and Head-UD are head rotated left-right and up-down respectively while walking. B: Top: Functional Gait assessment scores for all subjects, ordered in ascending age (65, 65, 67,71,79); data for mean aged subject groups adapted Walker 2007 for 70-79 (squares) and 80-90 (triangles). Bottom: Maximum lateral step placement for all 6 FGA trials that depend on lateral step. The horizontal dotted lines indicate the limit values for FGA scoring (see the text). C: Proposed additional measures of step width and movement duration, applicable to all trials along with limits for scoring shown in horizontal dotted lines (top and middle).

**Figure 5.** Experimental setup and validation experiment. A: Camera placement, with approximate distances relative to the left edge of the walkway. B: Experimental validation of step placements, using markers placed on the feet, sacrum and head. C: Validation of markerless data: step length, width and walking speed.

Variability in step placement across trials was qualitatively evident in overhead views of steps (Figure 4A top). We observed fluctuations for some FGA conditions in maximum lateral step placement (Figure 4A, bottom). For the tasks Normal, Speed Change, Head-RL and Head-UD, subjects exhibited qualitatively similar lateral displacement. For eyes open and eyes closed lateral displacements tended to be more variable. Compared to Normal, both Eyes Closed and Backwards walking demonstrated substantially larger displacements that were also more variable within subjects. Supplementary measure Step Width (Figure 4C) showed similar trends.

## Discussion

This study demonstrated the feasibility of a low-cost, AI-driven motion capture system to perform functional gait assessment. Through experiments on young and older adults, we showed that this system could accurately capture step placements and achieved a level of accuracy for step width and step length comparable to motion capture. Furthermore, the system’s utility was tested through Functional Gait Assessment (FGA) and provided suitable data comparable to previous data (Walker et al., 2007). The total cost of the three added consumer-grade web cameras and USB-extension cables was less than $200 CAD, and the analysis computer was inexpensive, meaning an existing computer at a clinic should be sufficient. Thus, modern open-source software appears ready to enable step-length, width and gait speed clinical testing for between 10% and 1% the cost of standard laboratory-based systems.

A critical question of markerless motion-capture systems is whether they are accurate enough for a given real-world application. The available open-source landmark “keypoint detection” inference models vary in 1) outputs: the set of points that are returned; 2) inputs the training set used to generate the model, in this case, human movements; and 3) details of model implementation, such as model architecture, training, and hyperparameters. Here we show that a system leveraging Google’s Mediapipe keypoint detection algorithm had sufficient foot position accuracy during stance phases to produce adequate step lengths and widths. Velocity information from head speed was also sufficient for walking tasks. Moreover, we expect the quality of available, open-source inference models to only improve, as new models continue to be developed worldwide (Chen et al., 2019; Nath et al., 2019). Thus, the potential for applications to health and industry will continue to broaden.

Functional Gait Assessment relies heavily on human visual identification of lateral foot placement (Figure 4). Step width is related but reflects the difference in lateral position of consecutive steps and its magnitude is thought to reflect control of lateral stability (Bruijn and van Dieën, 2018; Donelan et al., 2001; Kuo, 1999; Redfern and Schumann, 1994; Townsend, 1985; Wang and Srinivasan, 2014), given the instability of human gait in the frontal plane (Collins et al., 2005). Step length varies with walking speed (Grieve, 1968; Kuo, 2001) and gait speed decreases with age (Middleton et al., 2015). Continuous monitoring of both frontal and lateral steps may be particularly useful during unsteady walking conditions like accelerating, decelerating, or changing directions. Other stride-based techniques such as inertial measurement units are incapable of measuring step length and width, and thus have limited use for automatic scoring. Here we propose to use step width in addition to or in place of mediolateral foot placement (Figure 4C). It is widely used in the existing scientific literature (Donelan et al., 2001; Kuo, 1999; Redfern and Schumann, 1994; Townsend, 1985; Wang and Srinivasan, 2014), applies to non-straight walking paths, and thus is a more flexible measure of balance control.

Our system also yields simple automated assessments of gait speed. Gait speed is a crucial metric in fall risk assessment and traditional approaches often involve only average speed measurements over set distances, relying on manual timing by clinician. This process is repeated multiple times to observe deviations or deterioration (if any) in gait speed. Furthermore, gait speed is combined with other measures such as trunk or head acceleration variability or stride length variability and used to predict fall risk. Our system allows for a more detailed analysis by capturing speed trajectories over varying distances and conditions, such as those in FGA tests (particularly change in gait speed condition, Figure 4) as opposed to single speed measurement. Metrics like peak speed, mean speed, acceleration, and deceleration durations could be quantified from the trajectories. Therefore, our system can offer new metrics of speed for assessing gait speed changes and fall risk. Furthermore, these metrics can be compared with optimal control models (Carlisle and Kuo, 2023; Darici and Kuo, 2023, 2022) that predict speed and may provide testable mechanistic explanations underlying the difference between older and young adults. Furthermore, FGA is comprehensive, containing conditions involving turns and stepping over obstacles. Given our system is capable of capturing these movements, we presume that it can effectively be used for other simpler tests such timed up and go.

There are limitations to this study. The current system requires calibration to determine camera parameters via Direct Linear Transform that the software performs automatically. Calibration accuracy can affect results and takes between 5 and 10 minutes. Future applications will need to simplify calibration and hardware setup and provide feedback to the user that the system has been calibrated properly. We should note that if cameras are installed as permanent fixtures such as onto the ceiling of a room in a clinic, and if there is no potential for disturbances to the camera position and orientation, then calibration may need to be done only once. Another limitation is that statistical analysis between or within subjects was beyond the scope of this study, which was to demonstrate feasibility of gait assessment automation using open-source tools. For larger cohorts of patient populations these automated metrics such as movement duration (which replaces a clinician’s stopwatch), step width (which cannot readily be determined visually), and maximum lateral duration (which requires a labelled floor and clinician’s attention) may be used to track changes in behaviour across time, and possibly assist classification for fall risk.

One potential use for markerless motion capture is to integrate inertial measurement units (IMUs). Combining data from IMUs mounted on the feet (measuring linear acceleration and rotational velocity of the feet) can enhance the motion capture accuracy. Under normal conditions, movement can be measured with IMUs (Carlisle and Kuo, 2023; Darici and Kuo, 2022), but absolute orientation and location errors from accelerometer drift enlarge quickly in time without registration. Much larger capture volumes and perhaps multiple simultaneous subject recordings may also be realized with a combined setup. For instance, assuming two cameras are sufficient to record an area of 2 meters by 2 meters, 10 cameras might be integrated to cover a large real-world coordinate system and provide body position data across a large functional region. This setup may significantly improve the usage of IMUs and can facilitate various experiments to investigate how individuals navigate and control their step length, width, and speed in larger environments.

This study validated the feasibility of a low-cost motion capture system powered by computer vision and web cameras. With continued development, this system has the potential to transform clinical gait assessments by providing precise, automated measurements of key metrics such as step width and speed. It offers a cost-effective solution to reduce clinician workload, reduce measurement variability, and make advanced gait analysis accessible to a broader range of healthcare settings.

## Acknowledgments

This research was supported by NSERC Discovery (Wong), the University of Calgary Catalyst (Wong), and Biomedical Engineering Summer Student Scholarship (Cabak).

## Notes

### Competing Interest Statement

The authors have declared no competing interest.

### Summary of Updates

Paper rewrite given feedback from proofreaders; requested edits to primarily provide context with markerless studies, tighten writing.

